# Dysfunction of the colliculus-pulvinar pathway in children with developmental dyslexia

**DOI:** 10.64898/2026.06.20.733490

**Authors:** Yuzhu Ji, Yazhu Qian, Yue Wang, Junjun Li, Yizhen Li, Weiru Lin, Hong-Yan Bi, Peng Zhang

**Author notes:** Corresponding authors: Hong-Yan Bi; Peng Zhang. These authors contributed equally (Yuzhu Ji, Yazhu Qian).

## Abstract

While evidence suggests magnocellular deficits in the geniculostriate pathway in adults with dyslexia, neural deficits in the subcortical pathways during childhood remain unclear. Here, we used high-resolution fMRI to investigate subcortical abnormalities in Chinese children with developmental dyslexia. Fast achromatic motion stimuli and slowly drifting chromatic gratings were used to assess magnocellular (M) and parvocellular (P) functions, respectively. Relative to controls, children with dyslexia showed a selective reduction in responses to the M stimulus in the ventromedial pulvinar (vmPul) and the superficial layers of the superior colliculus (SCs), along with significantly reduced SCs-vmPul connectivity. Importantly, while vmPul responses to the M stimulus were positively associated with reading skills in healthy controls, this correlation was absent in children with dyslexia. Unlike previous findings in adults, the lateral geniculate nucleus (LGN) exhibited a non-selective reduction in responses to both stimuli, no volume reduction, and no correlation with reading ability. These findings demonstrate a selective deficit to achromatic motion processing in the colliculus-pulvinar pathway in children with dyslexia, which contributes to their reading difficulties. This early subcortical disruption differs from, and precedes, the neural deficits previously reported in the adult LGN, offering new insight into the developmental trajectory of dyslexia.

## Introduction

Developmental dyslexia (DD) is characterized by an impairment in the acquisition of reading skills despite normal intelligence, visual acuity, and education level (World Health Organization, 2022). In addition to the well-known phonological deficits (Ramus, 2003; Snowling, 2001), accumulating evidence suggests that many individuals with DD also exhibit impairments in visuosensory processing and visual attention (Stein, 2018), particularly those associated with the magnocellular (M) pathway and the dorsal visual stream (Stein, 1997). While these visual deficits could be important risk factors for dyslexia, direct functional evidence of subcortical deficits remain limited, especially in children.

Individuals with dyslexia have lower contrast sensitivity at low spatial and high temporal frequencies (Borsting et al., 1996; Cornelissen et al., 1998; Kevan & Pammer, 2008; Livingstone et al., 1991; Lovegrove et al., 1980, 1982; Martin & Lovegrove, 1984; Z.-L. Meng et al., 2022; Slaghuis & Ryan, 1999; J.-J. Wang et al., 2010), impaired motion perception (Demb et al., 1998; Qian & Bi, 2014; Wilmer et al., 2004), and deficits in attention (Facoetti, 2001; Facoetti et al., 2010; Fu et al., 2019; Goswami, 2015) and ocular motor controls (Bucci et al., 2008). A post-mortem study reported atrophy and disorganization of magnocellular, but not parvocellular (P) cells in the lateral geniculate nucleus (LGN) of the thalamus in a small sample (*n* = 5) of dyslexic brains (Livingstone et al., 1991). Recent MRI studies have shown that, compared with healthy controls, the left LGN of adults with dyslexia is smaller and differs in shape (Giraldo-Chica et al., 2015), exhibits weakened structural connectivity with the left middle temporal area (MT) (Müller-Axt et al., 2017), and shows abnormal lateralization of visual BOLD responses and T1-derived myelination in its M subdivision (Müller-Axt et al., 2025). Functional deficits have also been observed in the dorsal visual stream, including area MT and the posterior parietal cortex, both of which receive strong projections from the M pathway (Eden et al., 1996; Liu et al., 2022).

While subcortical visual deficits in DD have been studied primarily in the adult LGN, little is known about other subcortical visual nuclei, particularly the superior colliculus (SC) and the pulvinar, which play important roles in visuosensory processing, visual attention and ocular motor control. Visuosensory neurons in the superficial layers of the SC (SCs) receive direct retinal input from parasol (M-type) ganglion cells (Perry & Cowey, 1984) and cortical input from the early visual cortex, with similar response properties to neurons in the magnocellular layers of the LGN (Zhang et al., 2015). The deeper layers of the SC (SCd) receive input from frontoparietal attention networks and are involved in controlling attention and eye movements. The SC sends projections to the magnocellular portion of the visual thalamus, including the ventromedial pulvinar (vmPul) and the ventral LGN (Ghodrati et al., 2017; May, 2006), which in turn project to the dorsal visual stream (Arcaro et al., 2015; Kaas & Lyon, 2007). Compared to the vmPul, the ventrolateral pulvinar (vlPul) is more strongly interconnected with the early visual cortex and the ventral visual stream (Arcaro et al., 2015). The pulvinar has been suggested to play important roles in cortical computations, regulate information transmission between cortical areas, and contribute to visuosensory processing, perception and attention (Halassa & Kastner, 2017; Petersen et al., 1987). Increasing evidence suggests that SC and pulvinar are important parts of the language and reading network (Pugh et al., 2013; Zhang et al., 2021). Learning to read has also been shown to increase the functional connectivity between visual cortex and these subcortical nuclei (Skeide et al., 2017). Taken together, while these studies suggest that the SC and pulvinar may play important roles in learning to read, whether they exhibit functional abnormalities in DD remains unclear. Moreover, although subcortical deficits in DD have been studied almost exclusively in adults, the underlying pathology in children remains largely unknown. Given the extensive connections between the geniculo-striate and colliculus-pulvinar pathways, it remains unclear where the visual deficits in DD initially arise. Addressing this question would require investigating the functional abnormalities in the subcortical pathways of children with dyslexia, before long-term compensatory changes and reading experience in adulthood can obscure the origin of these deficits.

In the present study, we examined fMRI responses to M- and P-biased visual stimuli across subcortical and cortical visual areas in Chinese children with dyslexia. Compared to chronological age-matched controls (CA), children with DD showed a selective reduction in response to the M stimulus in the ventromedial pulvinar and the superficial SC, and a non-selective reduction to both stimuli in the LGN. Furthermore, the DD group exhibited reduced connectivity in response to the M stimulus in the SCs-vmPul and the LGN-hMT+ pathways.

## Results

To preferentially activate the P and M pathways (Derrington et al., 1984), visual stimuli were designed to differ in temporal frequency (low vs. high), and contrast (high-contrast chromatic vs. low-contrast achromatic stimuli). The P stimulus was an isoluminant high-contrast red-green grating drifting slowly at 0.5 Hz, presented in alternation with a yellow background (Fig. 1). The M stimulus was a low-contrast achromatic grating composed of concentric rings that expanded or contracted at 10 Hz, reversing motion direction every 4 seconds. Since P neurons respond to both chromatic and achromatic stimuli, the M stimulus was presented against the chromatic background to saturate the parvocellular activity, which can increase the selectivity for magnocellular activation. During the fMRI scan, participants were instructed to maintain central fixation and to detect occasional luminance changes of the fixation point. The performance of the fixation task was monitored online to ensure that all participants followed the task instruction.

**Figure 1.**
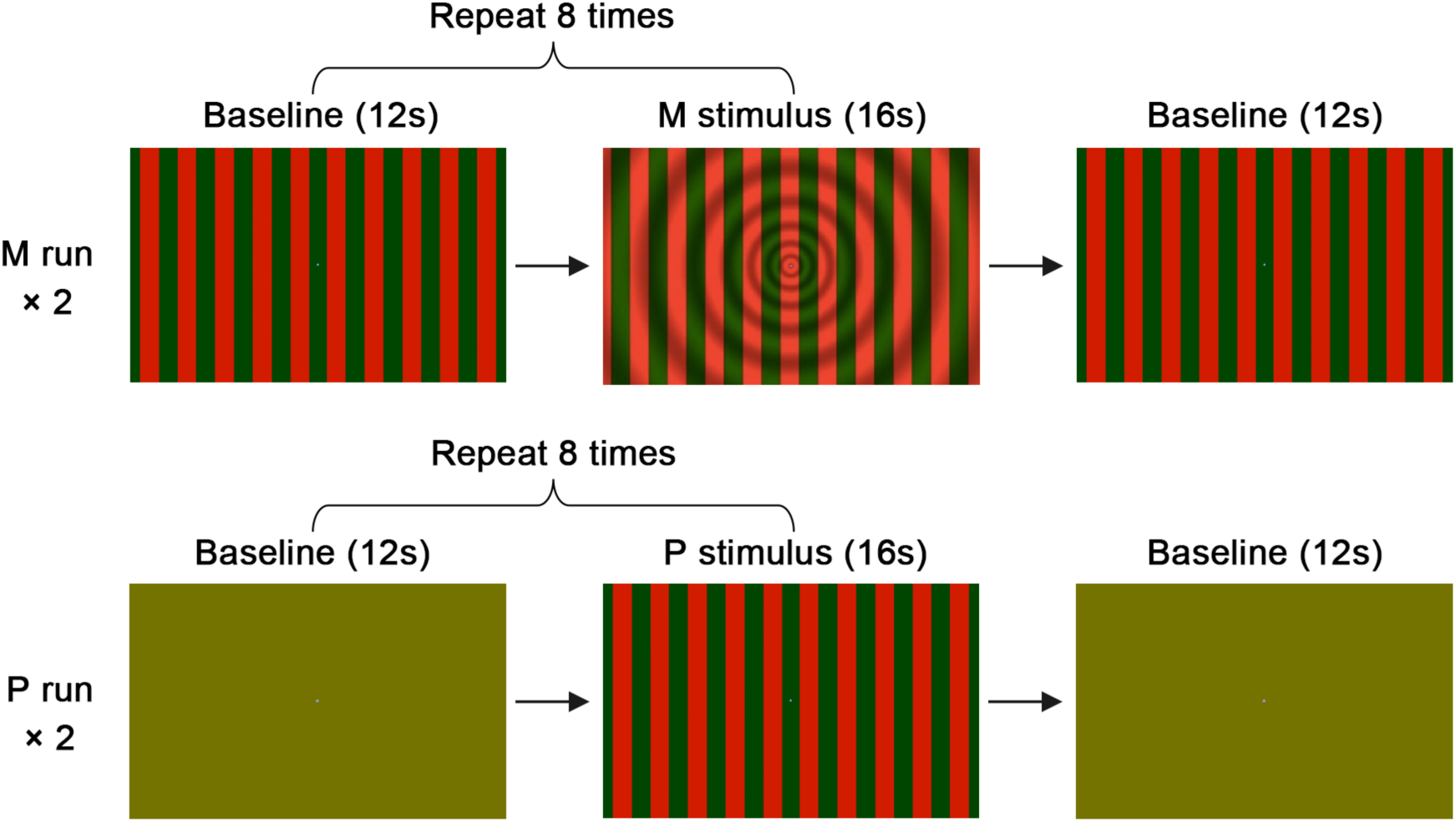
Stimuli and procedure for the fMRI experiment. In separate runs, M- and P-biased stimuli (16-s duration) were presented in alternation with baseline periods (12-s duration).

### Selective reduction of fMRI response to the M stimulus in the ventromedial pulvinar and the superficial SC

We first examined fMRI signals in the subcortical visual nuclei. Three-way repeated-measures (rm) ANOVAs were conducted for ROI-averaged responses in the LGN, pulvinar and SC (Fig. S4). The group (DD vs. CA) × stimulus (M vs. P) × hemisphere (left vs. right) interactions were not significant (all *ps* > 0.17), and no group × hemisphere interactions were observed (all *ps* > 0.31). Therefore, responses from the left and right hemispheres were averaged. Subsequently, a two-way rm-ANOVA with group and stimulus as within-subject factors was conducted for each region. Also, since there was no significant difference in M-selective deficits between males and females (all group × stimuli × sex interactions *ps* > 0.09), results were combined for both sexes.

#### LGN

As shown in the beta maps in Fig. 2a, M- and P-biased visual stimuli selectively activated the ventromedial and the dorsolateral portions of the LGN, respectively, consistent with the anatomical organization of the M (LGNm) and the P (LGNp) layers of the human LGN (Andrews et al., 1997; Hickey & Guillery, 1979; Zhang et al., 2015, 2016). Compared with the CA group, children with DD showed a reduction of fMRI responses to both M and P stimuli on the group-averaged beta maps (Fig. 2a, DD-CA). ROI-averaged responses in LGNp (Fig. 2b, right) showed a significant main effect of group (*F* (1,33) = 5.23, *p* = 0.029, 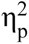 = 0.137), but no significant group × stimulus interaction (*F* (1,33) = 0.03, *p* = 0.865, 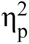 = 0.001), suggesting a non-selective reduction in responses to both stimuli. Pairwise comparisons further revealed a significantly reduced M response in DD compared to CA (*t* (33) = 2.523, *p* = 0.034, Holm corrected). Unlike previous findings in adults (Giraldo-Chica et al., 2015; Müller-Axt et al., 2025), our analysis revealed no significant between-group differences or group × stimulus interaction in LGNm responses (*ps* > 0.32). Furthermore, LGN volume did not differ significantly between the two groups in both hemispheres (*ps* > 0.72, Fig. S1).

**Figure 2.**
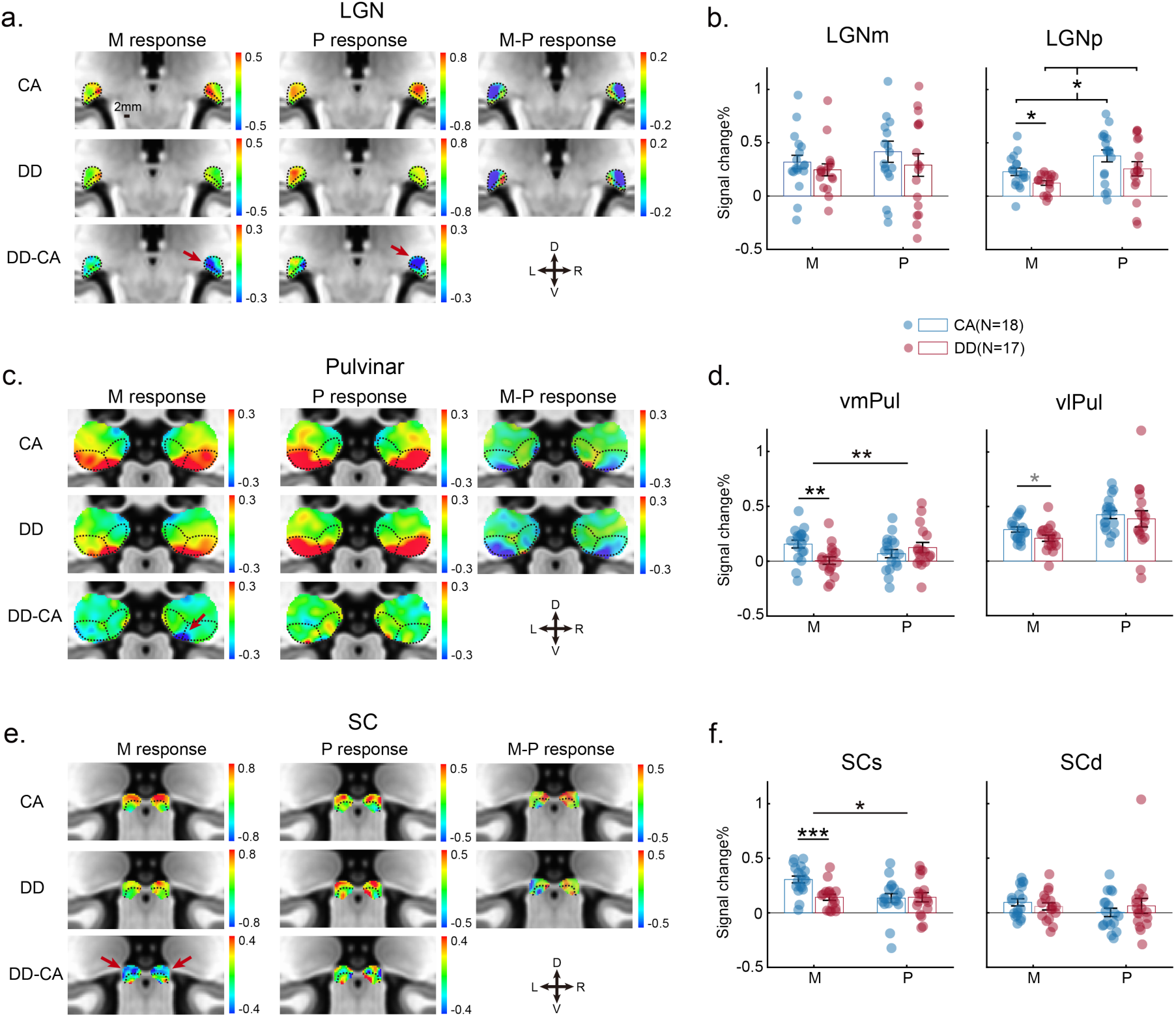
fMRI responses to M and P stimuli in subcortical visual nuclei. (a, c, e) Group-averaged beta maps for the CA group, DD group, and their difference (DD-CA) in the LGN, pulvinar and SC. M-biased activations (M-P > 0) were observed in the magnocellular layers of the LGN (LGNm), the ventromedial pulvinar (vmPul), and the superficial layers of the SC (SCs). In contrast, P-biased activations (M-P < 0) were located in the parvocellular layers of the LGN (LGNp) and the ventrolateral pulvinar (vlPul). Color bars represent percent change of BOLD signal from baseline. (b, d, f) ROI-averaged responses in functional subdivisions of the LGN (LGNm/LGNp), ventral pulvinar (vmPul/vlPul), and SC (SCs/SCd). Responses were averaged across hemispheres. Error bars indicate the standard error of the mean (SEM). Family-wise error: *p < 0.05, **p < 0.01, ***p < 0.001. The gray asterisk in vlPul denotes p < 0.05 before Holm correction.

#### Pulvinar

Group-averaged activation maps in Fig. 2c show that both M and P stimuli strongly activated the ventral pulvinar, consistent with its role in visuosensory processing and its intensive connections with the early visual cortices (Arcaro et al., 2015; DeSimone et al., 2015; Mai et al., 2015). Compared to the CA group, children with DD exhibited reduced activation to the M stimulus in the ventromedial portion of pulvinar (vmPul) (Fig. 2c, M response, DD-CA). Consistent with this, ROI-averaged fMRI response in the vmPul (Fig. 2d, left) showed a significant group × stimulus interaction (*F*(1,33) = 8.499, *p* = 0.006, 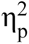 = 0.205). Follow-up comparisons showed that vmPul responses were significantly weaker in the DD group than in the CA group for the M stimulus (*t*(33) = 3.013, *p* = 0.005), but not for the P stimulus (*t*(33) = −0.927, *p* = 0.361). For the vlPul, no significant effect was observed. These findings suggest a selective reduction of fMRI response to the M stimulus in the ventromedial (magnocellular) pulvinar.

#### SC

Group-averaged activation maps (Fig. 2e) showed that M- and P-biased visual stimuli primarily activated the superficial layers of the superior colliculus (SCs). This observation is consistent with the established role of the SCs, which predominantly contains visuosensory neurons (May, 2006). Similar to the vmPul, SCs also showed a selective reduction in response to the M stimulus in DD compared to CA (Fig. 2e, M response, DD-CA). ROI-averaged responses in the SCs (Fig. 2f, left) revealed a significant group × stimulus interaction (*F*(1,33) = 4.82, *p* = 0.035, 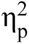 = 0.127). Furthermore, the DD group exhibited significantly weaker responses to the M stimulus (*t*(33) = 3.912, *p* < 0.001), but not to the P stimulus (*t*(33) = −0.134, *p* = 0.894). No significant group effect or related interaction were observed in the deep layers of the SC (SCd) (Fig. 2f, right, all *ps* > 0.334). These findings suggest a selective reduction of fMRI response to the M stimulus in DD compared to CA in the SCs, similar to those observed in the vmPul.

### Selective reduction of fMRI response to the M stimulus in hMT+

Figure 3a presents group-level activation maps on the cortical surface, with family-wise error controlled using the Threshold-Free Cluster Enhancement (TFCE) method (cluster *p* < 0.05). Relative to the CA group, the DD group exhibited significantly reduced responses to the M stimulus in the left V1, and the dorsal visual stream, as well as in the intraparietal sulcus (IPS) of the posterior parietal cortex bilaterally. However, a subsequent ROI-based analysis revealed no significant hemisphere × group × stimulus interactions in V1, hMT+, hV4, or IPS (all *ps* > 0.36). Accordingly, responses were averaged across hemispheres, and two-way repeated-measures ANOVAs with group and stimulus as within-subject factors were conducted for each region, following the same statistical procedure used for subcortical areas.

ROI-averaged responses revealed distinct patterns of response reduction across cortical regions (Fig. 3b). In hMT+, a selective reduction in response to the M stimulus was observed, as evidenced by a significant group × stimulus interaction (*F*(1,31) = 5.42, *p* = 0.027, 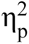 = 0.149) and a significant group difference specifically for the M stimulus (*p* < 0.001). V1 exhibited a significant main effect of group (*F*(1,31) = 9.07, *p* = 0.005, 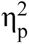 = 0.226), but no group × stimulus interaction (*F*(1,31) = 0.058, *p* = 0.811, 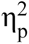 = 0.002). Pairwise comparisons revealed significantly reduced responses in the DD group for the M stimulus (*p* < 0.001, Holm-corrected), but not for the P stimulus (*p* > 0.1). In the IPS, a significant main effect of group was found (*F*(1,31) = 13.0, *p* = 0.001, 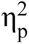 = 0.296), with the DD group showing significantly reduced responses to both the M (*p* = 0.006, Holm-corrected) and P (*p* = 0.008, Holm-corrected) stimuli compared to the CA group. No significant effect was observed in hV4. Together, these findings demonstrate a selective reduction of M responses in hMT+ and a non-selective reduction of fMRI responses to both stimuli in the V1 and IPS.

**Figure 3.**
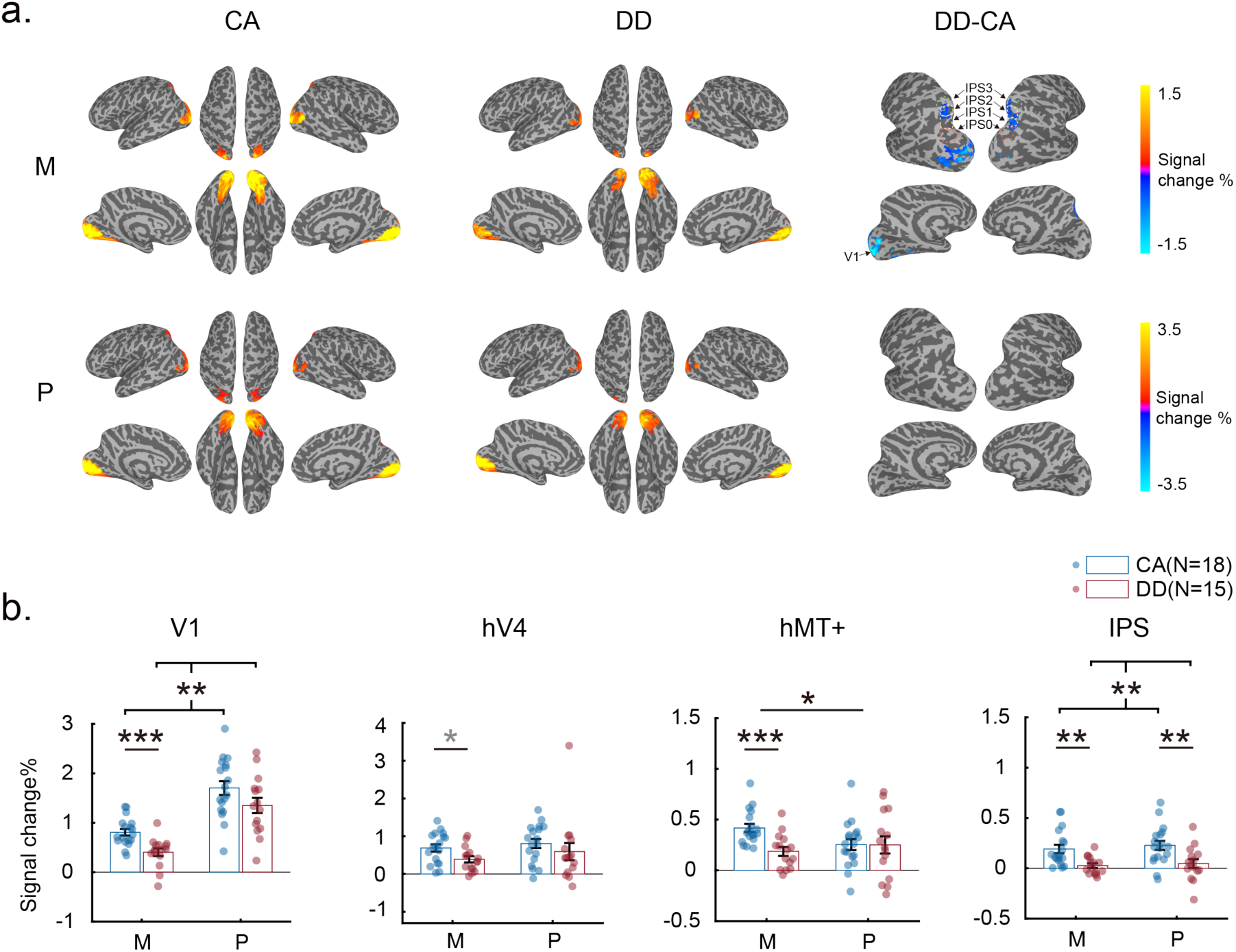
M and P responses in cortical visual areas. (a) Group-averaged activation maps on the cortical surface to the M and P stimuli for CA, DD, and DD-CA (TFCE controlled cluster *p* < 0.05). Statistical analyses were conducted within the retinotopic regions (Wang et al., 2015). Color bars denote percent change of BOLD response to the M and P stimuli. (b) Group-averaged responses to M and P stimuli in CA and DD groups, averaged across hemispheres. Error bars represent SEM. Red and blue dots represent data from individual participants. Family-wise error: \**p* < 0.05, \*\**p* < 0.01, \*\*\**p* < 0.001. The gray asterisk in hV4 indicates *p* < 0.05 before Holm correction.

### Reduced SCs-vmPul and LGNm-hMT+ connectivity in response to the M stimulus

To examine the causal influence of M signal loss among subcortical and cortical areas, we calculated effective connectivity using Dynamic Causal Modeling (DCM). Since this study focused on subcortical deficits, SCs, vmPul and LGNm were included in the analysis. For the cortical region, we selected hMT+, a representative area of the dorsal visual stream that exhibited selective M signal loss in our results (Fig. 3). The full DCM model in Fig. 4a was defined based on known anatomical connections (Arcaro et al., 2015; Ghodrati et al., 2017; Kaas & Lyon, 2007; May, 2006). In this model, LGNm, SCs and vmPul receive driving input from retina (green arrows). Fixed connections were defined between and within brain regions (black arrows). To investigate the M-specific connections, a modulatory effect was defined for the M stimulus condition on interregional connections (red dots). The DCM model was first fitted for each individual, and then the results entered a second-level Parametric Empirical Bayesian (PEB) analysis to assess the group-level differences between DD and CA (See Methods for details). For the group effect (DD vs. CA) on the M-modulatory connections, Bayesian model comparison indicated that the model with the highest posterior probability (model 48, Pp = 0.56) comprised connections from SCs to vmPul, from vmPul to LGNm, and from LGNm to hMT+ (Fig. 4a, right), which was consistent with the posterior connectivity pattern obtained by Bayesian model averaging (BMA) of the parameters over all possible models (Fig. 4b, right).

**Figure 4.**
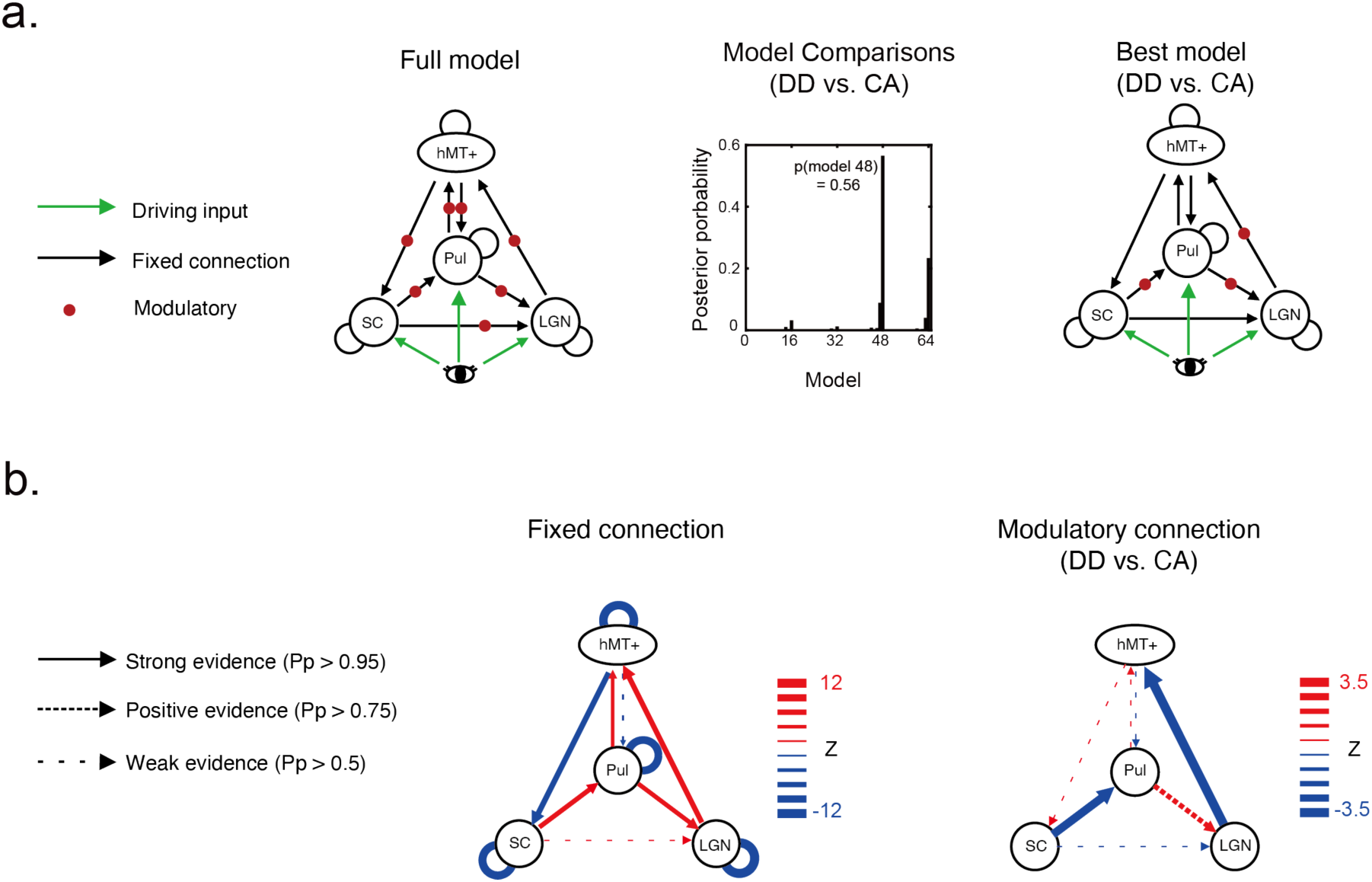
Effective connectivity results. (a) Left: The full DCM model illustrating the predefined connections among regions included in the analysis. Middle: Results of Bayesian model comparison testing the group-level differences (DD vs. CA) in the M stimulus-specific effects on connectivity. Right: The PEB model with the highest posterior probability. (b) Left: Bayesian model averaging (BMA) results of fixed connectivity (A matrix) across both DD and CA groups, including self-connections and between-region connections. Right: BMA results of the group differences (DD-CA) in the M-specific connectivity. Line width denotes the normalized strength (Z value) of functional coupling.

BMA results showed strong evidence (posterior probability, Pp > 0.95) for intrinsic connections among the subcortical nodes (SCs, vmPul, and LGNm) and the cortical area hMT+, as reflected in the fixed connectivity averaged across both DD and CA groups (Fig. 4b left). For the group-level differences on M-modulatory connections, there was strong evidence (Pp > 0.95) for reduced effective connectivity from the SCs to the vmPul and from the LGNm to the hMT+ in DD compared to CA (Fig. 4b right). These negative modulatory effects indicate that DD participants showed weaker transmission of M signals along both the colliculus-pulvinar pathway and the geniculo-extrastriate pathway. In contrast, a positive modulatory effect was observed for the vmPul-LGNm connection (Pp > 0.75), suggesting a potential compensatory influence within the subcortical network. Taken together, these results suggest that dyslexia is associated with reduced M-driven information transfer from subcortical structures to cortical motion processing areas.

### Ventromedial pulvinar responses to the M stimulus correlate with reading skills in CA but not in DD

Finally, to investigate whether the fMRI response to the M stimulus in the ventromedial pulvinar and the superficial SC can predict reading skills, we performed a correlation analysis within the CA and the DD groups separately. The fMRI response of vmPul to the M stimulus showed a significant positive association with a composite Z-score of key reading skills, including word reading, phonological awareness, and morphological awareness in the CA group (vmPul in Fig. 5, β = 0.541, *p_fdr_* < 0.001, adjusted R^2^ = 0.670). In contrast, no significant correlation was found in the DD group (vmPul, β = −0.117, *p_fdr_* = 0.494, adjusted R^2^ = −0.034). For both groups, no significant correlations were observed between reading scores and the M responses in the LGNm or SCs. A negative correlation was found between reading scores and P responses in the ventrolateral pulvinar (vlPul) in the DD group (Fig. S3), potentially reflecting a compensatory effect to the M deficits of vmPul.

**Figure 5.**
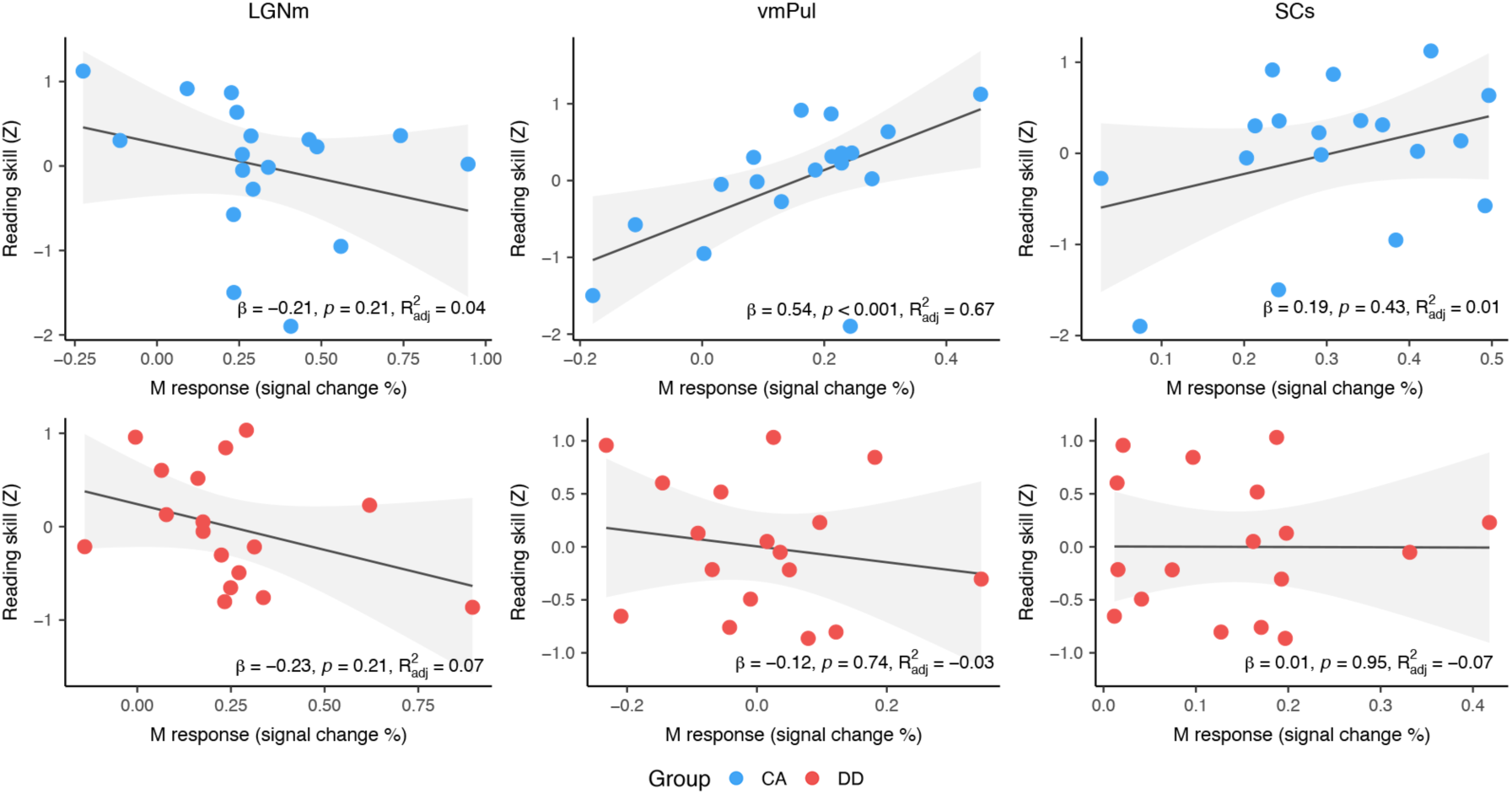
Associations between M responses in subcortical nuclei and reading skills. Associations were calculated by robust regression between M responses in LGNm, vmPul, and SCs, and the composite Z-score of word reading, phonological awareness, and morphological awareness within each group. The upper row (blue dots) and lower row (red dots) show the results for the CA group and the DD group, respectively. Each dot represents data from one participant. A significant positive association was observed between vmPul responses and reading skills in the CA group but not in the DD group. Shaded gray areas represent the 95% CIs of the fitted lines of robust regression.

## Discussion

Using M- and P-biased visual stimuli and high-resolution fMRI approach, this study investigated functional deficits in subcortical and cortical visual areas of Chinese children with developmental dyslexia (DD). Compared to chronological age-matched controls (CA), children with DD exhibited a selective reduction in response to the M stimulus in the ventromedial pulvinar and the superficial SC. Critically, while CA children showed a significant positive association between vmPul M responses and core reading skills, this relationship was absent in the DD group. Unlike previous findings in adults, LGN activity in children with DD showed a non-selective reduction to both M and P stimuli, with no significant correlation with reading abilities, and no differences in LGN volume. Finally, the DD group demonstrated reduced SCs-vmPul and LGNm-hMT+ connection in response to the M stimulus.

Basal ganglia and thalamic nuclei have been reported as important subcortical regions of the language and reading network (Nicolson & Fawcett, 2007; Wandell & Le, 2017). In a picture naming task, activation of the left thalamus was significantly correlated with reading scores in children beginning to read (Pugh et al., 2013). During character rhythming- and meaning-judgement tasks, bilateral thalamic nuclei and the left pallidum may act as network hubs in typically developing children, but not in children with DD (Zhang et al., 2021). Dysfunction of the medial geniculate nucleus, the auditory thalamus, has also been found in adults with dyslexia when performing a phonological task (Diaz et al., 2012). Here, we provide direct evidence that basic visual processing of fast achromatic motion is impaired in the vmPul and SCs in children with DD (Fig. 2). In typically developing children, vmPul responses to the M-biased stimulus were positively associated with reading skills, whereas this association was absent in the DD group (Fig. 5). These findings are consistent with previous studies showing magnocellular deficits in the LGN of adults with dyslexia, suggesting that subcortical visual deficits are present early in development. Given the similarity of the vmPul and SCs to the geniculate magnocellular pathways in term of their visual response properties and anatomical connections, our findings provide new subcortical evidence relevant to the magnocellular visual deficits in dyslexia (Stein, 1997, 2025).

The superficial layers of the SC receive magnocellular input from parasol ganglion cells of the retina (May, 2006). The ventromedial pulvinar receives input from the superficial SC and projects to the dorsal visual stream (Bridge et al., 2016; Kaas & Lyon, 2007); it also receives extensive retinal input in the developing primate brain (Mitchell & Leopold, 2015). The colliculus–pulvinar pathway represents an alternative, evolutionarily primitive route for processing transient visual information that operates in parallel with the geniculo-striate pathway. During early visual development in primates, the SC and pulvinar pathways might play a dominant role in processing behaviorally relevant information before maturation of the geniculate pathways and cortical networks (Bridge et al., 2016). This difference in the developmental trajectory is consistent with our observation that reading skills in typically developing children were associated with responses to transient achromatic motion in the ventromedial pulvinar, but not in the LGN (Fig. 5). In addition, there was no reduction in LGN volume (Fig. S1), in contrast to the shrinkage reported in adults with dyslexia (Giraldo-Chica et al., 2015; Livingstone et al., 1991). These findings suggest that in children learning to read, the colliculus-pulvinar pathway are more involved in processing transient information that supports the acquisition of reading skills. Early functional abnormalities in the SC and pulvinar may disrupt the development of the geniculate magnocellular pathway, which later exhibits functional and anatomical abnormalities associated with reading difficulties in adults with dyslexia (Giraldo-Chica et al., 2015; Livingstone et al., 1991; Müller-Axt et al., 2017, 2025). On the other hand, our results suggest impaired functional coupling in the magnocellular geniculo-extrastriate pathway (Fig. 4b right), which may also lead to magnocellular abnormalities within the LGN in the adult period.

Compared with the LGN, the pulvinar and the SC are more directly involved in higher cognitive functions, such as attention and ocular motor controls, in addition to low-level visuosensory processing of transient information. The pulvinar is an important subcortical region of the attention network, forming reciprocal connections with frontoparietal regions and visual cortices (Arcaro et al., 2015, 2018). Electrophysiological and lesion studies suggest a functional role of the pulvinar in attention control and regulating information transmission between cortical areas based on attentional demand (Saalmann et al., 2012; Saalmann & Kastner, 2011). The SC receives strong input not only from the early visual cortices, but also from the frontoparietal attention networks (May, 2006), enabling it to play a key role in orienting attention and eye gaze toward salient or goal-relevant information (Katyal et al., 2010; Schneider, 2011; White et al., 2017). Thus, functional deficits observed in the pulvinar and SC are highly consistent with the behavioral studies in DD showing deficits in attention (Facoetti, 2001; Facoetti et al., 2010; Fu et al., 2019; Goswami, 2015) and eye movement controls (Bucci et al., 2008), and in magnocellular processing such as motion detection and discrimination (Demb et al., 1998; Qian & Bi, 2014; Wilmer et al., 2004).

In cortical visual areas, our results indicate a tendency for magnocellular deficits to be more pronounced in the left hemisphere, particularly in the primary visual cortex (V1), and the dorsal visual stream (Fig. 3a upper). This finding is consistent with a left lateralized language network (Cohen et al., 2000; Hickok & Poeppel, 2007), and with the left lateralized functional abnormalities previously observed in adults with DD (Kronbichler et al., 2006; Müller-Axt et al., 2025). However, in the subcortical visual areas, we did not observe any significant lateralization effects. A number of reasons may explain this discrepancy. In children who are still learning to read, limited training and experience may result in weaker lateralization of the language and reading network compared to adults (Brown et al., 2005; Szaflarski et al., 2006; Turkeltaub et al., 2003), thereby exerting less influence on subcortical activity. In addition, given the lower signal-to-noise ratio in subcortical compared to cortical regions and the limited sample size of the present study, the absence of a lateralization effect in the subcortex may reflect the lack of sensitivity and detection power. The limited sample size may also explain the absence of a significant difference in M deficits between males and females. Finally, in contrast to the selective reduction of M responses in the pulvinar, SC and hMT+, a general signal reduction was observed to both M and P stimuli in the IPS. This non-selective signal reduction may reflect attentional deficits that broadly affected visual processing and learning ability.

In conclusion, we identified a selective deficit in processing transient achromatic motion within the colliculus–pulvinar pathway in children with developmental dyslexia. Unlike previous reports in adults, the LGN showed no selective response deficits or volumetric reductions. These findings suggest that visual dysfunctions in developmental dyslexia originate in the colliculus–pulvinar pathway during childhood, which may subsequently lead to the magnocellular deficits observed in the adult LGN. Together, these results provide new insight into the subcortical mechanisms and developmental trajectory of dyslexia, with implications for future research on diagnosis and intervention.

## Methods

### Participants

A total of thirty-five Chinese children participated in the fMRI study, including 17 children with developmental dyslexia (DD, 12 boys) and 18 chronological age-matched controls (CA, 11 boys). All participants were native Mandarin speakers and right-handed. Demographics and reading performance are shown in Table 1. Written informed consent was obtained from each participant’s guardian before participating the experiment. This study was approved by the ethics committee of the Institute of Psychology, Chinese Academy of Sciences.

**Table 1.** Demographics and reading performance of CA and DD.

| | **CA (*n* = 18)** | **DD (*n* = 17)** | *t*/$\chi^2$ | *p* | *Cohen's d* |
| --- | --- | --- | --- | --- | --- |
|  | *Mean (SD)* | *Mean (SD)* |  |  |  |
| Age (years) | 10.40 (0.69) | 10.04 (0.67) | 1.55 | 0.13 | 0.53 |
| Sex (male/female) | 11/7 | 12/5 | 0.35 | 0.56 |  |
| IQ (standard score) | 115.22 (13.18) | 108.65 (10.61) | 1.62 | 0.12 | 0.55 |
| Chinese Character Recognition (items) | 2904.35 (416.02) | 1870.76 (479.41) | 6.82 | < **0.001** | 2.31 |
| Word reading (items) | 105.61 (13.98) | 80.82 (13.69) | 5.30 | < **0.001** | 1.79 |
| Phonological awareness (items) | 28.56 (1.65) | 24.12 (3.46) | 4.88 | < **0.001** | 1.65 |
| Morphological awareness (items) | 16.56 (2.12) | 13.53 (1.81) | 4.53 | < **0.001** | 1.53 |

We estimated the sample size using G*Power (Faul et al., 2007). Based on a previous study related to visual thalamic connectivity in dyslexia (Müller-Axt et al., 2017), which reported a group × cortical area interaction effect of η² = 0.07 (equivalent to f = 0.274), power analysis indicated that 30 participants would provide 80% power at α = 0.05. Therefore, we recruited 35 participants to ensure adequate statistical power.

We screened 1,120 children in grades 3 to 5 from three primary schools in Beijing to identify DD based on the following criteria: (1) scoring at least 1.25 standard deviations below the grade mean on the Chinese Character Recognition Test (CCRT; X. L. Wang & Tao, 1993), a widely used standardized vocabulary test for screening dyslexia in Mandarin-speaking children (Feng et al., 2020; X. Meng et al., 2011; Tian et al., 2024; J.-J. Wang et al., 2010; Yang et al., 2022); (2) a nonverbal intelligence quotient (IQ) above 85, measured by the Raven’s Standard Progressive Matrices (Raven et al., 1996); (3) normal or corrected-to-normal vision and hearing; and (4) no history of attention deficit hyperactivity disorder (ADHD) or other neurological disorders. Similar procedures for diagnosing dyslexia have been adopted in previous studies in mainland China (e.g., Feng et al., 2020; Yang et al., 2022).

Controls were defined based on the following criteria: (1) IQ-, sex-, and age-matched to the DD group; (2) scores on the CCRT were within or above the average range for the same grade level (> −0.5 SD); and (3) normal or corrected-to-normal vision and hearing, with no history of neurological disorders.

All participants completed the reading-related skill tests, including word reading, phonological awareness and morphological awareness. Independent two-sample *t*-tests revealed that children with dyslexia performed significantly worse than controls on all reading measures. See Table 1 for more details about participants.

### Stimuli and Procedures

Visual stimuli were generated in MATLAB (Mathworks Inc.) with psychophysics toolbox extension (Brainard, 1997; Pelli, 1997) on a MacBook pro computer, presented by a MRI safe projector at 1024×768 pixel resolution at 60Hz. The color look-up table of the projector was calibrated to have a linear luminance output.

Figure 1 shows the experimental procedures and visual stimuli during the fMRI scan. The M stimulus consisted of sine wave concentric rings at 30% luminance contrast, expanding or contracting at 10 Hz, with the spatial frequencies that decreased with eccentricity. The spatial frequency *f* (in cycles per degree, cpd) as a function of eccentricity *θ* (in degrees) was given by:

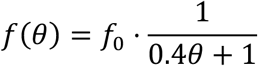

Where *f*_0_ is the spatial frequency at the stimulus center (*f*_0_ ≈ 1.5 cpd), and *f* dropped to about 0.2 cpd at the outer edge of the stimulus (*θ* ≈ 16.45^∘^).

The P stimulus was an isoluminant red/green grating with the spatial frequency of 0.25 cpd, moving from left to right or vice versa at 0.5 Hz. The red/green isoluminance point was determined by experienced adult observers using a minimal flicker procedure, as pilot testing indicated that the task was too challenging for children to perform reliably. This approach ensured a stable and consistent stimulus parameter across all participants. The temporal frequency for the M stimulus (10 Hz) and the P stimulus (0.5 Hz) was selected to preferentially engage magnocellular and parvocellular pathways, respectively, consistent with established physiological characteristics of these pathways (e.g., Derrington et al., 1984; Merigan & Maunsell, 1993).

To isolate neural responses biased toward the M and P pathways, we employed two separate functional runs. In P runs, the P stimulus was presented in 16-second blocks, interleaved with 12-second fixation baselines. In M runs, the P stimulus served as a continuous baseline, onto which the M stimulus was superimposed. This design aimed to saturate P neuron responses, thereby enhancing the selectivity of M responses. As shown in Fig. 2a, M and P stimuli preferentially activated the ventromedial and dorsolateral portion of the LGN, respectively, consistent with the anatomical locations of the M and P layers. Moreover, M and P stimuli also selectively activated hMT+ and hV4 in the dorsal and ventral visual streams (Fig. 3b, CA group), further supporting the validity of the stimulus design. Four runs were scanned for each participant, consisting of two runs each for the M or P stimulus. Each run contains 8 stimulus blocks and 9 baseline blocks. During fMRI scans, participants were instructed to maintain fixation and press a button to occasional color changes of the fixation point. Four runs of similar stimuli has been shown to provide sufficient statistical power to detect robust M and P activations in the LGN (Wen et al., 2021; Zhang et al., 2015, 2016).

### MRI data acquisition

MRI data were collected with a 3T scanner (MAGNETOM Prisma^fit^, Siemens, Erlangen, Germany) at the Beijing MRI Center for Brain Research. A gradient echo planar imaging sequence was used to acquire functional images (2-mm isotropic voxels, 54 axial slices of 2 mm thickness, 96× 96 matrix, TR/TE = 2000/31.8 ms, flip angle = 80°, multiband factor = 2, no parallel imaging). High-resolution anatomic volume was obtained with a T1w MPRAGE sequence (1-mm isotropic voxels, 192 sagittal slices of 1-mm thickness, 258 × 258 matrix, TR/TE = 2600/3.03 ms, flip angle = 8°).

### MRI data preprocessing

Preprocessing of functional data was done in AFNI (Cox, 1996), including the following steps: slice timing correction, EPI distortion correction with nonlinear warping, rigid body motion correction, alignment of corrected EPI images to the T1-weighted anatomical volume (cost function: lpc), and per-run scaling as percent signal change. To minimize image blur, all spatial transformations were combined together and applied to the functional images in one interpolation step. General linear models with a fixed HRF (Block4 in AFNI) were used to estimate BOLD signal change from baseline for each stimulus condition. Motion parameters were included as regressors of no interest.

### Subcortical ROIs and volume-based analysis

To compare LGN volume between CA and DD groups, anatomical regions of interest (ROIs) of the LGN were manually delineated on each participant’s T1w MPRAGE images, where the LGN appears darker than surrounding tissue. Delineation was performed by an experienced researcher (Y.Q.) who was blind to the group identity. To validate the reliability of the manual delineation, the manually drawn LGN ROIs were transformed into MNI space and overlaid to generate a probability map (Fig. S2a), which showed that the ROIs were clearly separable from the pulvinar and largely overlapped with the LGN region in MNI space.

Subcortical data were normalized to the symmetric MNI template of the CIT168 atlas (Pauli et al., 2018) using nonlinear warping in Advanced Normalization Tools (ANTs; Avants et al., 2011). To improve registration accuracy in subcortical ROIs, we applied a spherical mask centered on the visual thalamus and upper brainstem. The normalized data were then resampled to 0.6 mm isotropic resolution. Functional subdivisions within these subcortical nuclei were subsequently defined based on known anatomical characteristics and functional atlases.

A normalized layer index map was generated on the LGN template. Two layers of voxels were defined corresponding to the ventral and dorsal surfaces of the LGN. For the rest of voxels, we calculated a layer index as the ratio of the shortest distances to the dorsal and ventral surfaces of the LGN (0 to 1 correspond to ventral and dorsal surfaces, respectively). The M and P ROIs were determined from the layer index map according the volume ratio of M/P layers of human LGNs (M/P = 1:4; Fig. S2b; Andrews et al., 1997).

The pulvinar was parcellated according to the Barron et al. (2015) atlas, focusing on two ventral (or visual) subdivisions: the ventromedial pulvinar (vmPul) and the ventrolateral pulvinar (vlPul). Because the ventral pulvinar is relatively large and has a low signal-to-noise ratio, we further refined the vmPul and vlPul ROIs using voxel selection. Specifically, a leave-one-subject-out (LOSO) procedure was applied to identify voxels showing significant activation to the M and P stimuli (M+P > 0, *p* < 0.05) for the left-out participant. This approach ensured independent voxel selection thus avoided the risk of double dipping. Consistent with this subdivision, the M-P contrast revealed stronger negative responses in the vlPul (Fig. 2c), supporting the functional relevance of the anatomical parcellation.

To delineate the superficial and deep layer compartments of the SC, two layers of voxels were first defined from the superficial and deep surfaces of the SC. Then a normalized depth map was calculated for each voxel as the ratio of the shortest distances to the superficial and deep surfaces (0 and 1 correspond to the superficial and deep surfaces, respectively). Finally, the volume of the SC was split into a superficial (SCs) and a deep (SCd) layer compartments at a normalized depth of 0.5 (Fig. S2c).

### Cortical ROIs and surface-based analysis

Cortical surfaces were reconstructed from T1w anatomical volume for each participant using FreeSurfer. Surface data were analyzed in AFNI/SUMA (Argall et al., 2006) and in SPM. Data from 15 children with DD and 18 controls were used for the surface-based analysis due to surface reconstruction failure in 2 DD participants. Statistical volumes from the preprocessing analysis were projected to the cortical surface and were spatially smoothed with a 4-mm FWHM Gaussian kernel, followed group-level random effects analyses. Group-level activations on the cortical surface were corrected for multiple comparisons using non-parametric permutation test in CoSMoMVPA (Oosterhof et al., 2016) in Matlab, and Threshold-Free Cluster Enhancement (TFCE) was used to control family-wise error of clusters (Spisák et al., 2019). Retinotopic ROIs of cortical visual areas were defined on the cortical surface using the probabilistic atlas of Wang et al. (2015), and then mapped into the native volume space of each individual. The IPS was treated as a single ROI including all subregions (IPS0–IPS5). Due to the limited scan time of children participants, we did not measure an independent functional localizer for hMT+. Instead, hMT+ was defined by combining the atlas regions TO1 and TO2 in the lateral temporal cortex along the middle temporal gyrus, consistent with previous reports on the location of hMT+ (Amano et al., 2009).

### Dynamic causal modeling

The DD group exhibited a selective deficit in response to the M stimulus, particularly in vmPul, SCs and hMT+. To further investigate the information flow among these regions, we performed a dynamic causal modeling (DCM) analysis. The LGNm, SCs, vmPul and hMT+ were included as the M-biased ROIs. ROI-averaged time series from the M runs were extracted for each region and concatenated across runs for individual-level model estimation. The full DCM model is shown in Fig. 4a. The M stimulus was modeled as exogenous driving inputs (matrix C), entering the system exclusively through the LGNm, SCs and vmPul. Fixed connection (matrix A) included both inter- and intra-regional connections, and was specified based on known anatomical connections among these regions (Bridge et al., 2016; Ghodrati et al., 2017; Kaas & Lyon, 2007; May, 2006). Modulatory effects (matrix B) were specified to test M stimulus-specific influence on inter-regional connectivity.

Subject-level models were fitted using the variational Bayesian scheme in SPM12 (release 2020-Jan-13). Group-level effects were estimated using the parametric empirical Bayes (PEB) framework. The second-level design matrix included a constant term and a binary group regressor (0 for CA, 1 for DD). Bayesian model reduction (BMR) and Bayesian model comparison (BMC) was used to identify the most parsimonious model, and Bayesian model averaging (BMA) was applied to generate posterior estimates while accounting for model uncertainty. To better visualize the effective connectivity, we transformed the posterior estimates of the modulatory parameters into standardized Z values by dividing the posterior mean (Ep) by the posterior standard deviation (derived from the diagonal of the posterior covariance matrix).

### Statistics

A three-way repeated-measures (rm) ANOVA was first employed on the ROI-averaged responses in each brain region, with stimuli (M vs. P) and hemisphere (left vs. right) as the within-subject factors, and group (DD vs. CA) as the between-subject factor. Since the statistics of rm-ANOVA is not sensitive to violations of normality (Blanca et al., 2017, 2023), we didn’t remove outliers in the parametric analysis. To assess associations between brain activations and reading skills, robust linear regression models were fitted in each group. Given the relatively small sample size and the potential influence of outliers, robust regression was adopted to reduce sensitivity to extreme observations and to provide more stable parameter estimates. Continuous variables were z-standardized prior to analysis, and standardized regression coefficients (β) are reported. P values were corrected for multiple comparisons across ROIs using the false discovery rate (FDR) procedure. Group-level effective connectivity was assessed using a parametric empirical Bayes (PEB) framework. First, a PEB model was estimated across all participants ([PEB, RCM] = spm_dcm_peb(GCM_all, M)). Model comparison and reduction were then performed using Bayesian model comparison ([BMA, BMR] = spm_dcm_peb_bmc(PEB)), which applies Bayesian model reduction (BMR) to evaluate nested models without re-estimation. To account for model uncertainty, Bayesian model averaging (BMA) was applied, and inferences were based on the posterior parameter estimates and their associated posterior probabilities obtained from the BMA results.

Two DD participants were excluded from the cortical surface analysis due to failed surface reconstruction. As a result, there were 33 participants in the cortical analyses.

## Data and code availability

Data and code to reproduce the main findings of this study will be made available upon publication. Raw MRI data cannot be made publicly available since sharing these data is not covered by the ethics clearance.

## Acknowledgments

This study was supported by Brain Science and Brain-like Intelligence Technology -National Science and Technology Major Project (2021ZD0200500 to H.-Y.B., 2022ZD0211900 and 2021ZD0204200 to P.Z.), and the National Natural Science Foundation of China [32371119] to H.-Y. B. We are deeply grateful to the children and their parents who participated in this study.

## Author contributions

Y.J., H.-Y.B., and P.Z. conceived the research; Y.J. recruited participants and collected the behavioral data; Y.J., and Y.W. collected the MRI data; W.L., and P.Z. programmed the visual stimuli; Y.J., Y.Q, Y.W., J.L., and Y.L. analyzed the data; Y.J., Y.Q, and P.Z. wrote the draft of the manuscript; Y.J., Y.Q, H.-Y.B., and P.Z. revised the manuscript.

## Competing interests

The authors declare no competing interests.

## Notes

### Competing Interest Statement

The authors have declared no competing interest.

