## Supplementary methods and figures for "Dysfunction of the colliculus-pulvinar pathway in children with developmental dyslexia"

### Supplementary Information

#### Measures of reading skills

*Chinese Character Recognition Test.* This is a widely used standardized vocabulary test for screening Mandarin-speaking Chinese children with dyslexia (Feng et al., 2020; X. Meng et al., 2011; Tian et al., 2024; Wang et al., 2010; Yang et al., 2022). In this test, participants were instructed to write down a compound word using each target morpheme character. For children in grades 3, 4, and 5, the test included 206, 174, and 210 characters of varying difficulty, respectively. The task of making words rather than naming characters was adopted because Chinese contains many homophones, and correct pronunciation alone does not ensure understanding of character meaning.

*Word Reading.* In this task, a list of 172 Chinese characters of increasing recognition difficulty (i.e., more strokes and lower word frequency) was presented on paper. Participants were asked to read each character aloud in sequence without a time limit. The number of correctly read characters was recorded as the test score. The reliability of this task (Cronbach's alpha) was 0.88 (Z.-L. Meng et al., 2022).

*Phonological awareness.* This test employed the oddball paradigm (Bradley & Bryant, 1978). On each trial, three single Chinese characters were presented auditorily through headphones, and participants were asked to select the item that was phonologically odd. Each participant completed 10 trials for each of three oddity types: onset, rime, and lexical tone (e.g., /meng3/ differed from /gao1/ and /bao4/ in rime). The number of correct responses was recorded as the test score. The reliability of this test (Cronbach's alpha) was 0.75 (Z.-L. Meng et al., 2022).

*Morphological awareness.* In this task, two bimorphemic words sharing one morpheme were presented on each trial (e.g., 信 [letter or believe] in 信封 [envelope] and 信任 [trust]). Participants were asked to judge whether the shared morpheme carried the same meaning in both words. The task consisted of 20 trials, and the number of correct responses was recorded as the test score. The reliability of this task (Cronbach's alpha) was 0.75 (Z.-L. Meng et al., 2022).

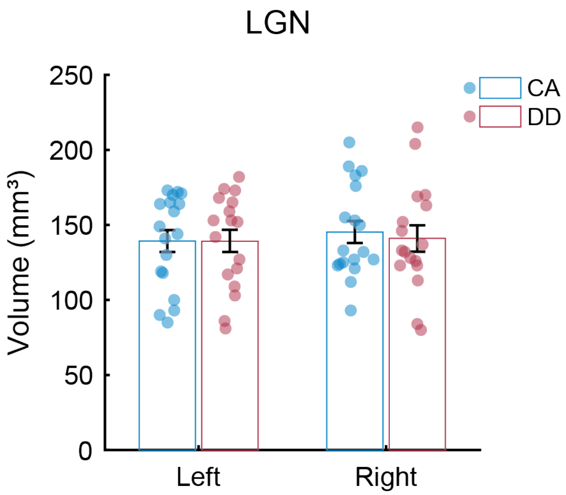


**Figure S1. LGN volume comparison between CA and DD groups.** To assess whether the LGN structural alterations observed in dyslexic adults are present in childhood, we compared LGN volumes between groups. No significant group differences were found in either the left or right LGNs (both *ps* > 0.72). Error bars represent SEM.

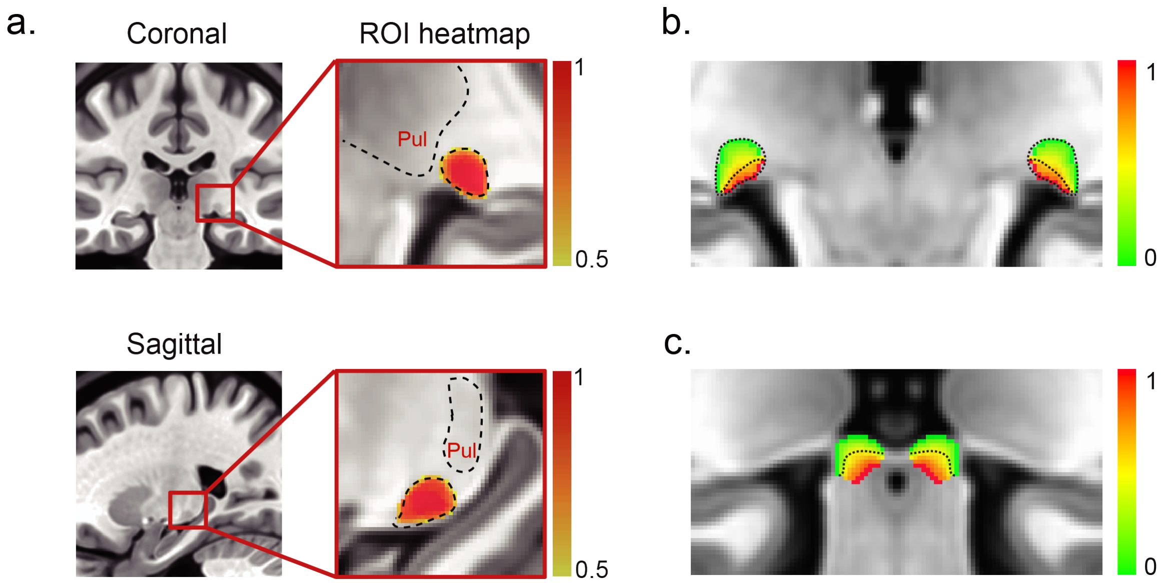


**Figure S2. Anatomical boundaries and layer segmentations of the LGN and the SC.**

(a) Black dotted lines on the coronal and sagittal views illustrate the manually delineated anatomical boundaries of the LGN and the pulvinar on the symmetric MNI template of the CIT168 atlas. To assess the accuracy of spatial normalization to the MNI template, the manually defined LGN ROIs in native space were transformed to MNI space. The colorbar indicates the overlap of manually defined LGN ROIs across participants (0.5 and 1 indicate 50% and 100% overlap, respectively). The map shows good registration quality, with the ROIs falling within the anatomical boundary of the LGN.

(b) Normalized layer index map and ROIs for the M and P layers of the LGN. Two layers of voxels were first defined corresponding to the ventral and dorsal surfaces of the LGN. For the remaining voxels, we calculated a layer index as the ratio of the shortest distances to the dorsal and ventral surfaces (0 and 1 correspond to the dorsal and ventral surfaces, respectively). The M and P ROIs were then determined from the layer index map according to the volume ratio of M to P layers in the human LGN (1:4). Black dotted lines denote the boundary of the LGN and the boundary between the M and P ROIs.

(c) Normalized depth map and superficial and deep layer ROIs for the SC. Two layers of voxels were first defined from the superficial and deep surfaces of the SC. A normalized depth map was then calculated for each voxel as the ratio of the shortest distances to the superficial and deep surfaces (0 and 1 correspond to the superficial and deep surfaces, respectively). The volume of the SC was split into a superficial (SCs) and a deep (SCd) layer compartment at a normalized depth of 0.5. Black dotted lines denote the boundary between SCs and SCd.

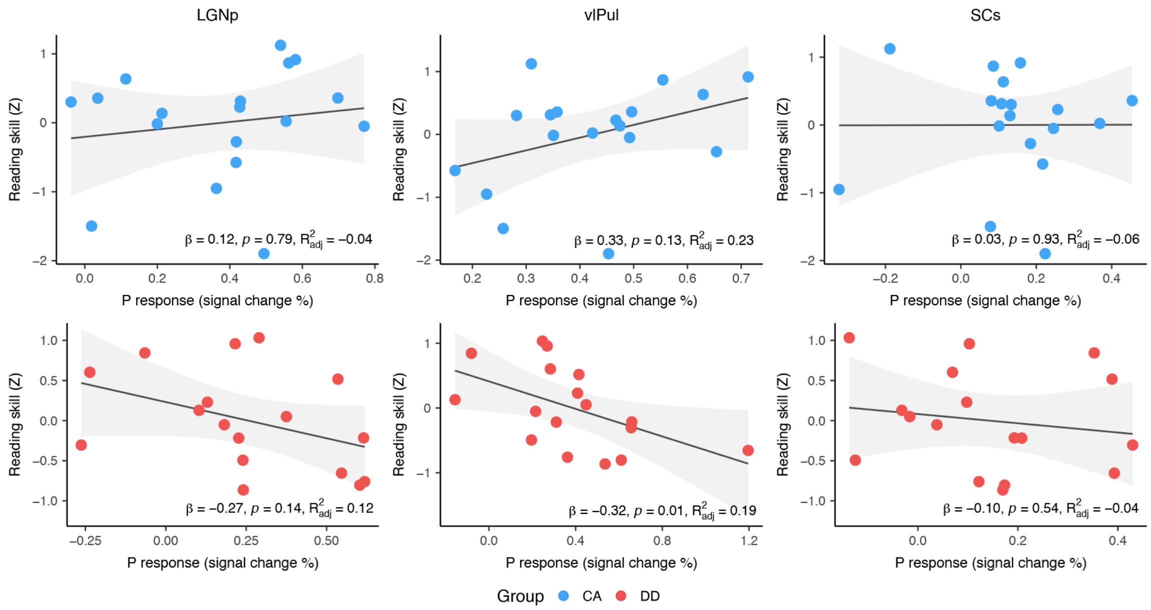


**Figure S3. Associations between P responses in subcortical nuclei and reading skills.** Associations were calculated by robust regression between P responses in LGNp, vlPul, and SCs, and the composite Z-score of word reading, phonological awareness, and morphological awareness within each group. The upper row (blue dots) and lower row (red dots) show the results for the CA group and the DD group, respectively. Each dot represents data from one participant. In the DD group, vlPul activation showed a significant negative association with reading skills. Shaded gray areas represent the 95% CIs of the fitted lines of robust regression.

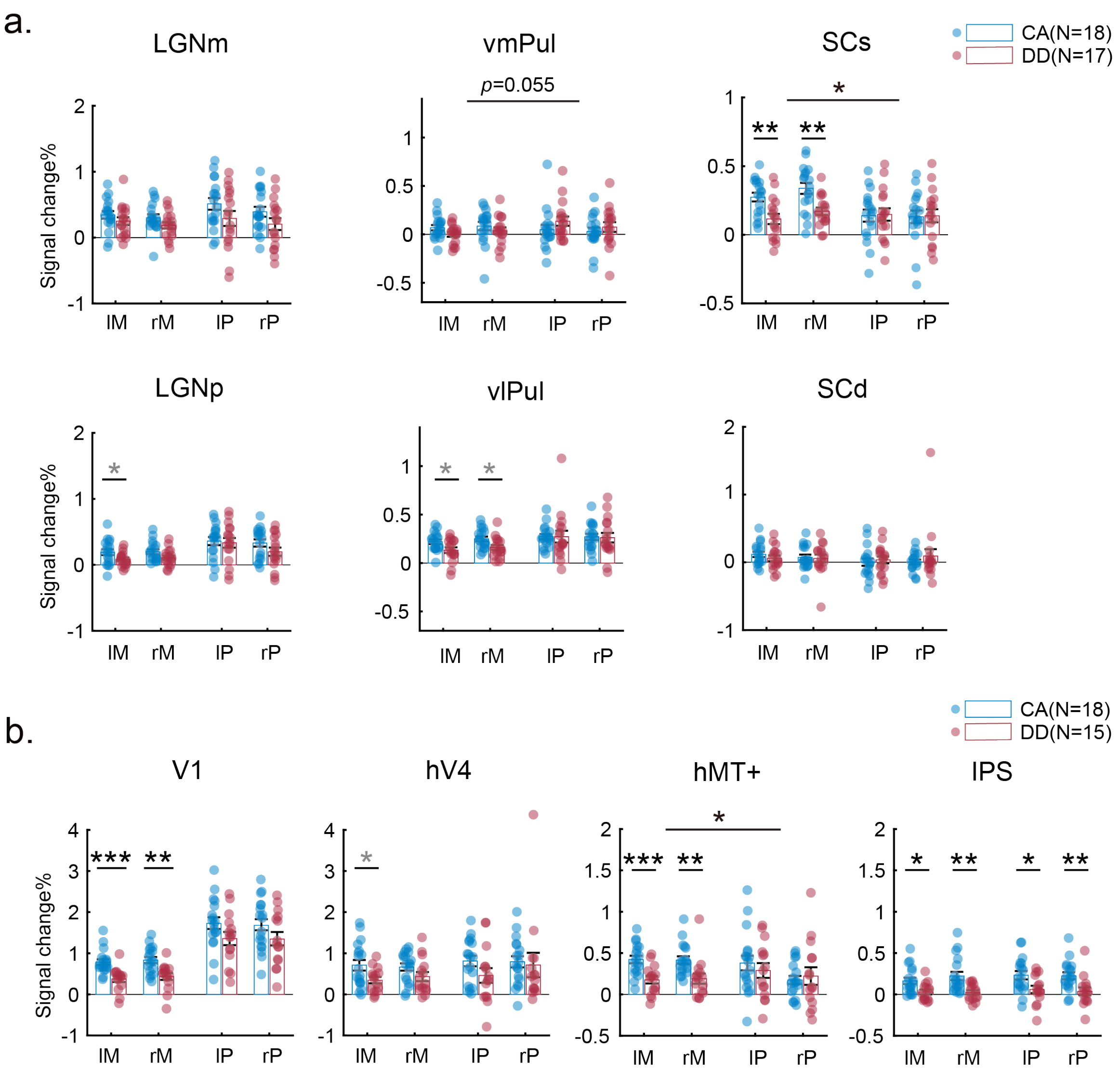


**Figure S4. ROI-averaged responses to M and P stimuli separately for the left and the right hemispheres.** The M-selective deficits were similar in the left and the right hemisphere of subcortical regions. While the activation map on the cortical surface shows a M-selective deficits lateralized to the left hemisphere (Fig. 3a), no significant lateralization effects were found in the ROI-averaged responses of cortical regions. Error bars represent SEM. *$p < 0.05$, **$p < 0.01$, ***$p < 0.001$. The gray asterisks denote $p < 0.05$ before Holm correction. The p value and * above a long black line indicate the statistical significance of group × stimulus interaction.
